# Increased atherosclerotic plaque and anti-oxidized low density lipoprotein in anti-Ro60 SLE autoantibody subset

**DOI:** 10.1101/2023.04.04.535634

**Authors:** Biji T. Kurien, James Fesmire, Swapan K. Nath, R. Hal Scofield

## Abstract

**Objective:** Premature atherosclerosis is associated with systemic lupus erythematosus (SLE). We previously showed an association of anti-Ro60/La/Ro52 to anti-oxidized LDL in SLE. Here, we hypothesized that atherosclerotic plaque will be associated with anti-oxidized LDL (anti-oxLDL)/anti-lipoprotein lipase (ALPL) in a specific SLE autoantibody sub-set (anti-Ro60 positive, anti-RNP positive or extractable nuclear antigen antibody negative).

**Methods:** We carried out a case-control study (one time-point testing) of plaque, ALPL, anti-oxLDL, anti-low density lipoprotein (ALDL) or anti-LDL in 114 SLE and 117 age/sex matched controls. Total cholesterol, LDL, HDL, triglycerides, and HDL-Trig were also measured. Students t test was used for statistical analysis

**Results:** Interestingly, plaque was highest in SLE subset with anti-Ro60 (23/114). Plaque and anti-oxLDL were statistically significantly elevated in the anti-Ro60 SLE subset (1.3 +/-1.66, p<0.01; 0.26 +/-0.16, p<0.002 respectively) compared to controls (0.54 +/-1.26; 0.165 +/-0.13 respectively), but not anti-LPL/anti-LDL. Plaque was significantly elevated (0.9 +/-1.71; p< 0.05) in the SLE subset without anti-ENA (63/114) compared to controls. Other antibodies in this subset were not statistically different compared to other SLE subsets or controls. Only anti-oxLDL was significantly elevated (0.29 +/-0.27; p<0.005) in SLE subset with anti-RNP (14/114) compared to controls, while none were elevated in anti-SmRNP subset (6/114). We did not find significant differences in lipids between various SLE subsets.

**Conclusion:** Plaque segregates in anti-Ro and ENA negative groups either with or without anti-oxLDL. It will be clinically important if cardiovascular events are augmented in SLE anti-Ro subset having elevated anti-oxidized LDL antibodies.

**Key points:**

1. Plaque and anti-oxLDL significantly elevated in anti-Ro60 SLE subset
2. Only plaque significantly elevated in SLE subset without antibodies against extractable nuclear antigens
3. It will be clinically important to see if augmented cardiovascular events occur in SLE anti-Ro subset having elevated anti-oxidized LDL antibodies.

## Introduction

Premature atherosclerosis is an important late complication of systemic lupus erythematosus (SLE), while being a significant issue also in patients with early SLE. There is increased prevalence of atherosclerotic plaque formation in SLE subjects, as well as elevated risk of cardiovascular disease (CVD).^1-4^ Non-traditional risk factors such as cytokines, chemokines, autoantibodies as well as traditional risk factors contribute to the development of CVD.^5,6^ Of late it has become evident that atherosclerosis and its corollary, CVD, is an inflammatory ailment and that the immune system influences the development of disease.^7^ It is therefore of interest to investigate the elevated risk of CVD in SLE since immune mechanisms in human atherosclerosis could be elucidated. Autoantibodies targeting oxidized LDL, cardiolipin and β2 glycoprotein 1 are associated with SLE and anti-phospholipid syndrome related vasculopathies. Immunochemical epitopes found on oxidized-LDL are seen in atherosclerotic lesions.^8,9^ Antibodies to oxidized LDL are increased in atherosclerosis and are also found in SLE, diabetes, hypertension, and pre-eclampsia.^8-11^

Autoantibodies in SLE target a 60,000 molecular weight protein (Ro60 or SS-A) or 48,000 molecular weight La (SSB) autoantigen of the Ro RNP particle, associated non-covalently with one or more of four short uridine-rich human cytoplasmic RNAs (hY RNAs). Up to 50% of SLE subjects have anti-Ro60. Anti-La occurs in considerably fewer SLE subjects.^12,13^ SLE autoantibodies also target the autoantigen Ro52 (also known as TRIM21).^14,15^

Sm and nuclear ribonucleoprotein (nRNP) antigens are also commonly targeted in SLE. These proteins are involved in the splicing of pre-mRNA in association with U small nuclear RNAs. Anti-Sm autoantibodies are found in the sera of roughly 20 to 25% of all SLE subjects. These antibodies form a part of the criteria for SLE classification and are highly specific for SLE.^13^

It is important to note that mortality from lupus manifestations has diminished consequent to better treatment modalities. However, deaths due to CVD from atherosclerosis in SLE have not. CVD actually is responsible for more than a third of all deaths in SLE subjects.^16,17^

Ultrasound instrument has been of great use to detect atherosclerotic plaque and also to measure carotid artery intimamedia thickness (IMT).^18^ Owing to ease of visualization and reproducibility, IMT is preferably measured in the common carotid artery.^19,20^ Furthermore, internal carotid artery measurement has been used successfully to detect and measure carotid plaque in subjects with atherosclerosis/cardiovascular-related conditions.^21,22^

Our recently reported work showed increased vulnerability of SLE subjects with anti-Ro 60, La and Ro 52 to develop anti-oxidized LDL.^23^ In this study we tested the hypothesis that atherosclerotic plaque will be associated with anti-oxidized LDL and anti-lipoprotein lipase in a specific autoantibody subset of SLE.

## Subjects and methods

### Subjects

Data collected from an earlier study ^11^ was analyzed after obtaining study approval from the Institutional Review Board of the Oklahoma Medical Research Foundation. This study used 114 SLE subjects (104 women and 10 men) and 117 age/sex matched controls. The subjects were not on any lipid lowering medication. The study subjects met SLE 1982 revised classification criteria of the American College of Rheumatology. OMRF Clinical Immunology Laboratory, a CLIA approved facility carried out the serological studies. ANA was tested by indirect imunofluorescence using a HEp-2 substrate. Anti-double stranded DNA was determined by *Crithidia lucilliae* immunofluorescence and autoantibodies to extractable nuclear antigens by double immunodiffusion. Normal controls were selected to match the SLE subjects for sex and age. None of the controls were taking any lipid lowering medications. The study was approved by the Oklahoma Medical Research Foundation Institutional Review Board.

## Methods

All assays were carried out in the previous study and reported ^11^, and are described briefly here.

### Carotid bilateral ultrasound

Carotid bilateral ultrasound (USBL) was performed by the Cardiovascular Section, Department of Medicine, University of Oklahoma Health Sciences Center. The plaque scores were measured on a scale from 0 to 10. The study group was provided a duplex carotid screen (both arteries) by means of Doppler sonography. The atherosclerotic plaque burden is expressed as a sum of values determined in both arteries. Subjects only with a value entry in the ultrasound test were taken into account for statistical analysis.^11^

### Anti-LPL, anti-oxidized LDL or anti-LDL ELISA

The assays were carried out in the previous study.^11^ Lipoprotein lipase, oxidized LDL or LDL was coated on ELISA plates, blocked with milk and subject sera was added at a 100-fold dilution and incubated overnight. The plates were washed, incubated with anti-human IgG alkaline phosphatase conjugate followed by substrate and O.D. read at 405 nm.

### Lipid determination and HDL/LDL isolation

Cholesterol and triglycerides determination as well as HDL/LDL isolation was carried as described.^11^

### Statistical analysis

Values are presented as means ± standard deviation (SD). Statistical analysis was carried out using Student’s t-test with *p* <0.05 considered as statistically significant.

## Results

A previous study from our ^11^ group investigated the extent of coronary risk due to anti-lipoprotein lipase and anti-oxidized LDL in the context of carotid plaques in SLE subjects and normal controls. The study found anti-lipoprotein lipase (anti-LPL) associated with oxidatively modified LDL, production of anti-oxidized LDL antibody, plaque formation and coronary risk in some SLE patients.^11^

Since we recently showed that there is increased susceptibility of SLE subjects with anti-Ro 60, La and Ro 52 to have anti-oxidized LDL and that immunization with Ro60 autoantigen induces anti-oxidized LDL (ox-LDL) antibodies (unpublished data), we tested the hypothesis that anti-lipoprotein lipase antibodies and anti-oxidized LDL will be associated with atherosclerotic plaque in a specific autoantibody subset of SLE.

Double immunodiffusion studies showed that sixty-three SLE subjects (55.26%) did not have autoantibodies against extractable nuclear antigen (ENA), 14 (12.2%) had antibodies against ribonucleoprotein (RNP), 23 (20.18%) had anti-Ro60, 6 (5.26%) had anti-SmRNP, 4 (3.5%) had unidentified precipitin lines and 4 (3.5%) had miscellaneous antibodies. Plaque was highest in the SLE subset with anti-Ro60 autoantibodies compared to all other lupus subsets and normal controls (Table 1, Figure 1). While elevated compared to other SLE subsets, the plaque values were not statistically different. However, plaque in this subset was statistically different compared to plaque in the normal controls (p<0.01) (Table 1, Figure 1). Antibodies against oxidized LDL was also elevated in this subset compared to other SLE sub-sets. However, anti-oxidized LDL antibodies in this subset was only significant compared to normal controls (p<0.002) (Table 1; Figure 2).

**Table 1:**
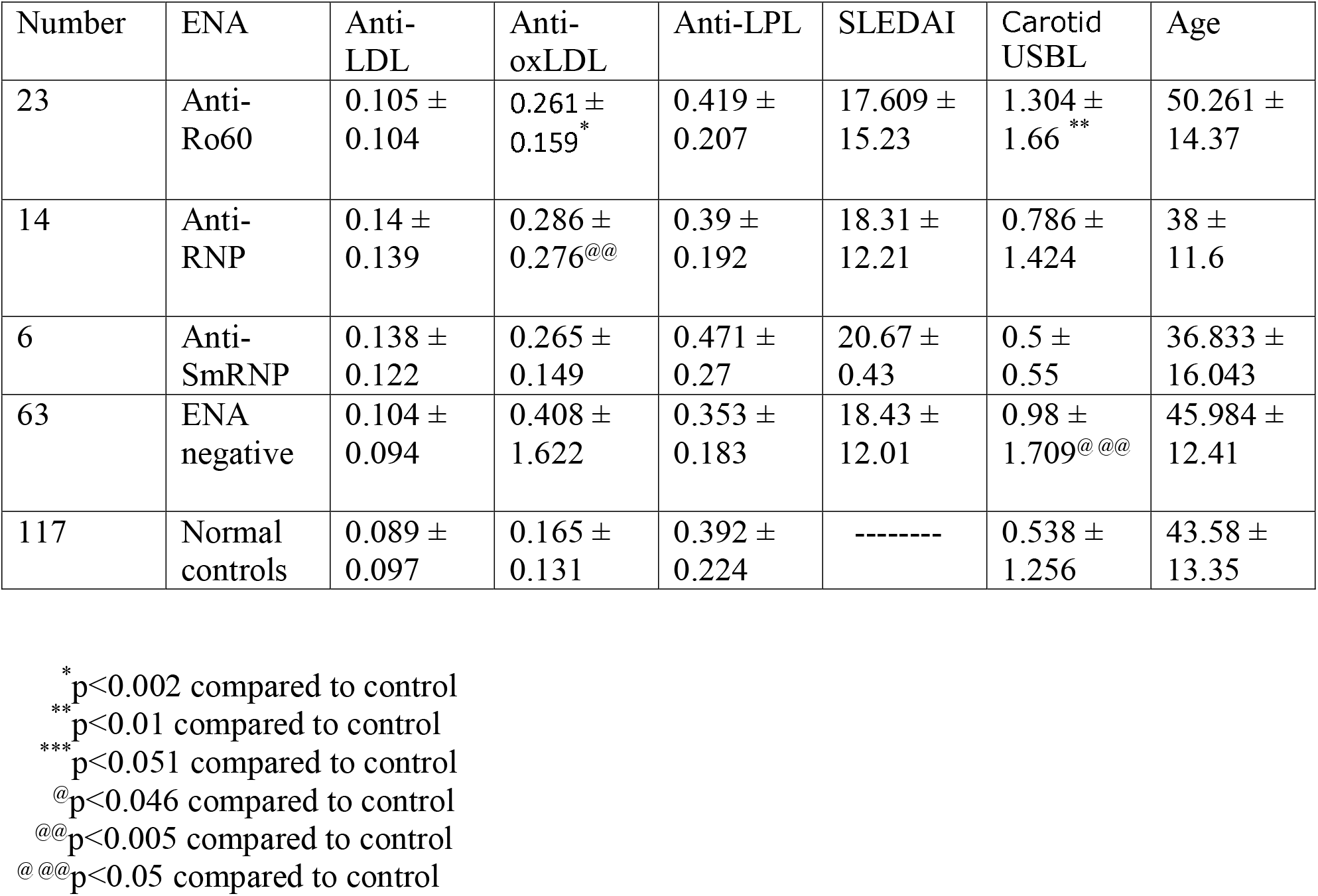
Antibodies to low density lipoprotein (LDL), oxidized LDL (oxLDL) and lipoprotein lipase (LPL) as well as SLE disease activity index (SLEDAI), carotid bilateral ultrasound (USBL) in subsets of lupus patients and controls.

**Figure 1:**
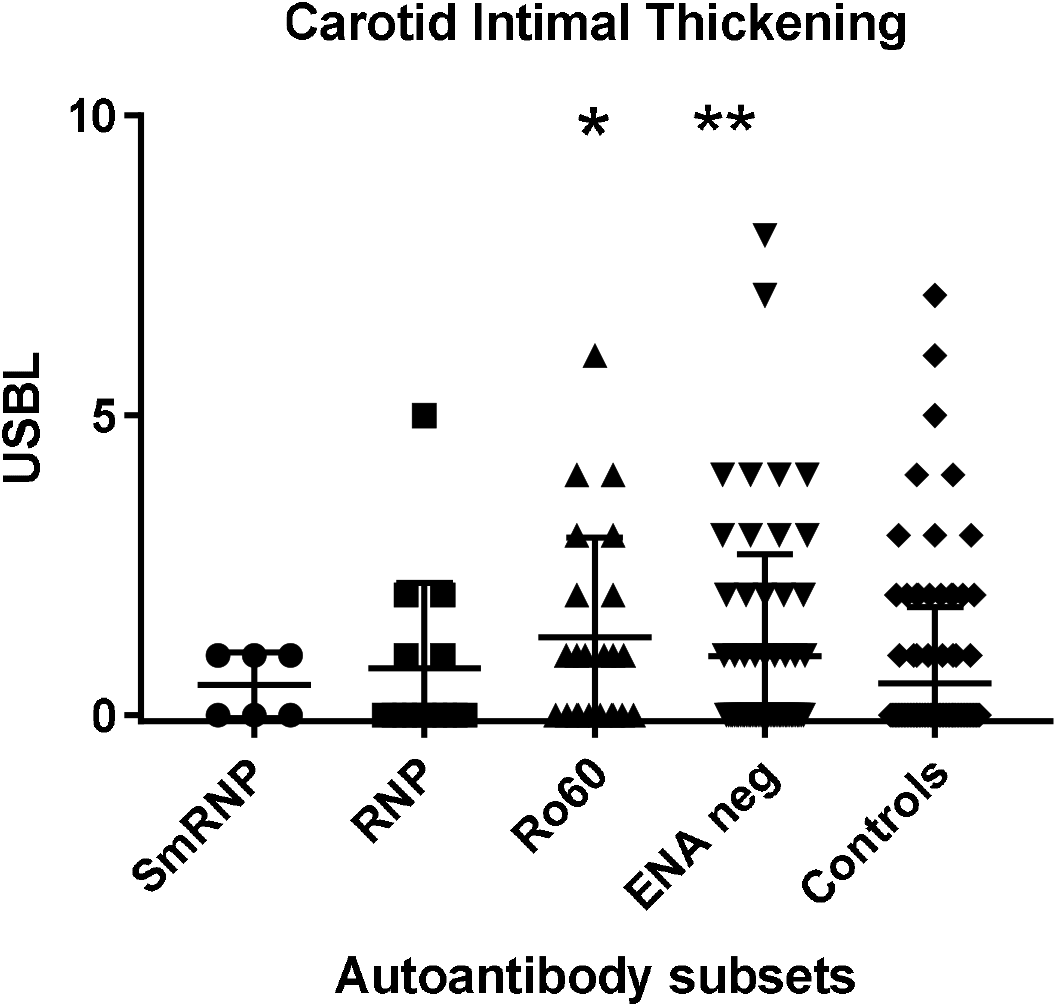
Carotid intimal thickening in SLE autoantibody subsets and normal controls. Carotid bilateral ultrasound (USBL) was used to determine intimal thickening and plaques scores expressed on a scale from 0-10. SLE subjects were divided into subsets based on their autoantibody profile determined by immunodiffusion studies and analyzed for USBL scores. ^*^p<0.01 compared to control ^**^p<0.05 compared to control

**Figure 2:**
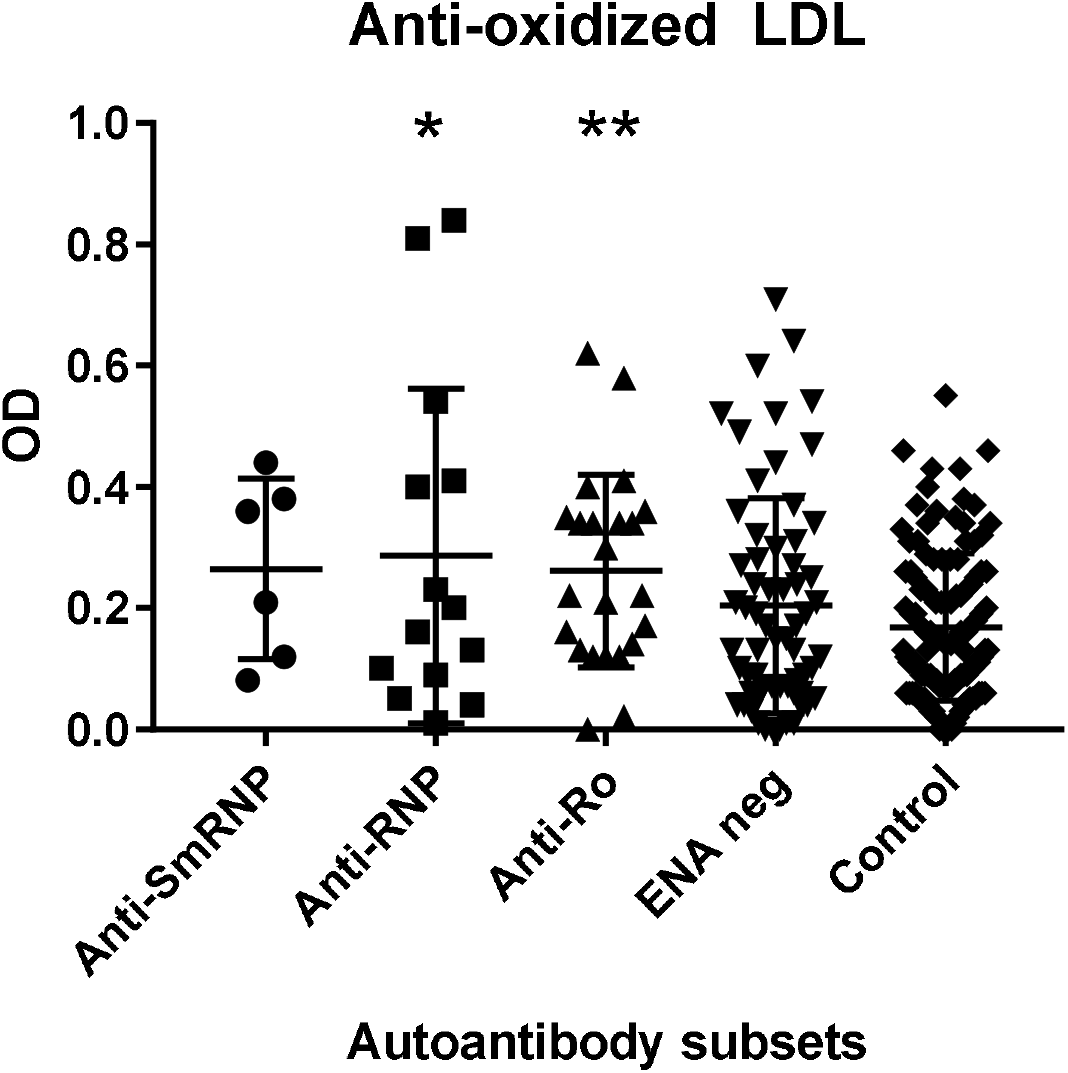
Anti-oxidized LDL antibodies in SLE autoantibody subsets and normal controls. Antibodies against oxidized LDL was determined as mentioned in ‘Subjects and methods’. SLE subjects were divided into subsets based on their autoantibody profile determined by immunodiffusion studies and analyzed for anti-oxidized LDL antibodies. ^*^p<0.005 compared to control ^**^p<0.002 compared to control

Plaque was lower in the SLE subset that did not have autoantibodies against extractable nuclear antigens compared to the SLE subset with anti-Ro60. However, the difference was not statistically significant. Interestingly, plaque was significantly elevated in this subset compared to normal controls (p< 0.05). Antibodies against oxidized LDL, LPL and LDL was not statistically different in this sub-set compared to the other SLE sub-sets as well as normal controls (Tables 1,2; Figures 1-3).

**Table 2:** Total cholesterol, total triglycerides, low density lipoprotein (LDL)-cholesterol, and high density lipoprotein (HDL)-cholesterol in subsets of lupus patients and normal controls.

| # | ENA | Total cholesterol | Total tri-glycerides | LDL-cholesterol | HDL-cholesterol |
| --- | --- | --- | --- | --- | --- |
| 23 | Anti-Ro60 | 179.92 ± 48.68 | 123.38 ± 72.3 | 101.54 ± 38.73 | 53.16 ± 17.46 |
| 14 | Anti-RNP | 176.04 ± 38.56 | 166.56 ± 99.05<sup>*</sup> | 101.86 ± 99.52 | 47.72 ± 10.61 |
| 6 | Anti-SmRNP | 188.15 ± 52.42 | 136.55 ± 74.06 | 116.1 ± 37.33 | 45.68 ± 9.62 |
| 63 | Negative | 212.05 ± 56.04<sup>***</sup> | 170.95 ± 111.54<sup>**</sup> | 124.12 ± 40.26 | 55.16 ± 23.05 |
| 117 | Normal controls | 192.31 ± 39.48 | 117.89 ± 69.61 | 109.46 ± 29.91 | 55.34 ± 15.09 |

<sup>\*</sup>p<0.025 compared to control <sup>\*\*</sup>p<0.0002 compared to control <sup>\*\*\*</sup>p<0.007 compared to control

**Figure 3:**
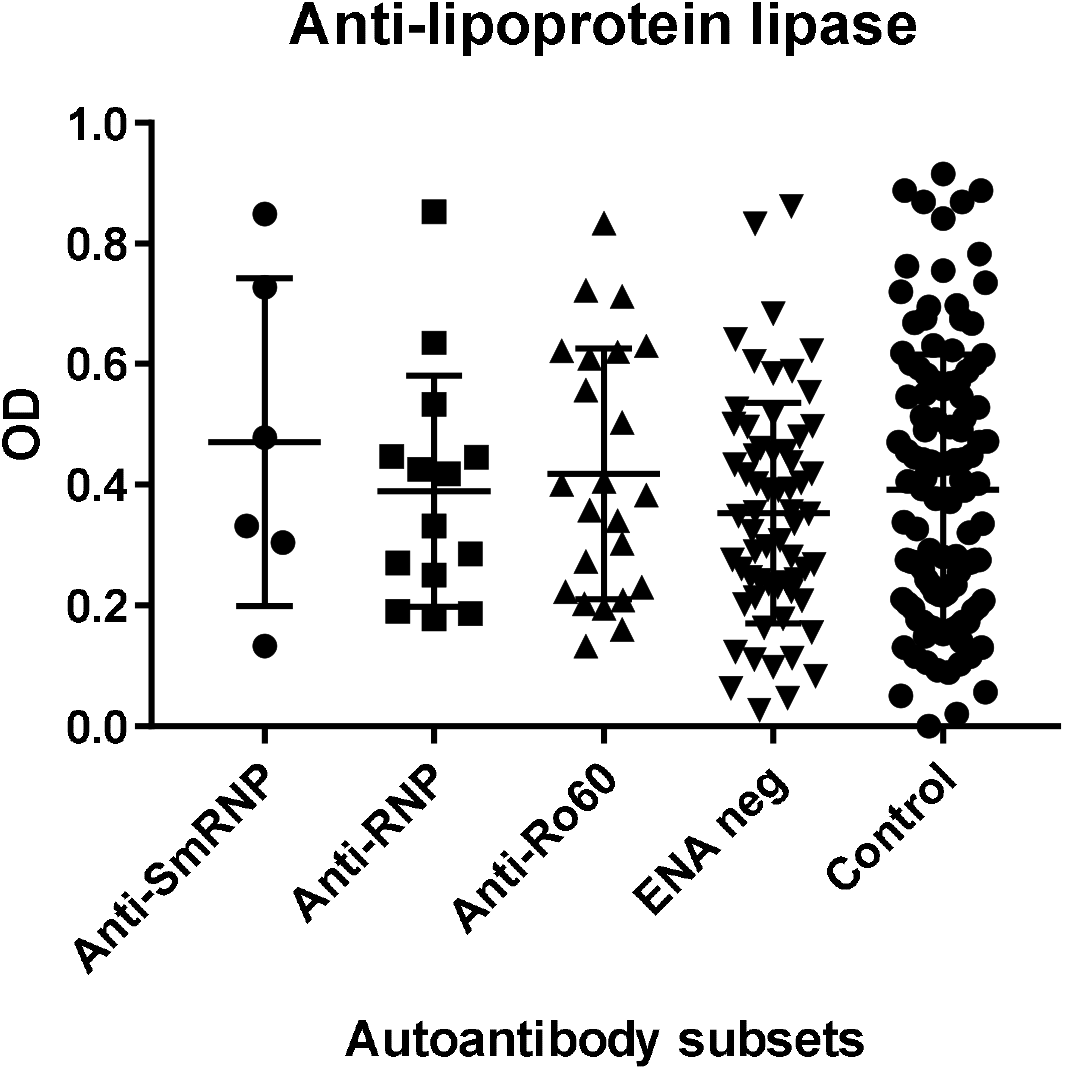
Anti-lipoprotein lipase antibodies in SLE autoantibody subsets and normal controls. Antibodies against lipoprotein lipase was determined as mentioned in ‘Subjects and methods’. SLE subjects were divided into subsets based on their autoantibody profile (determined by immunodiffusion studies) and analyzed for anti-lipoprotein lipase antibodies.

The SLE subset with anti-RNP antibodies had significantly higher levels of anti-oxidized LDL antibodies (p<0.005) compared to control group, but not within other SLE subsets (Table1, Figure 2). Plaque and antibodies against LPL or LDL were not different in this subset compared to control group as well as other SLE subsets (Table 1, Figure 1). The anti-SmRNP SLE subset did not behave significantly different from normal controls and other SLE subsets with respect to plaque as well as antibodies directed against oxidized LDL, LDL or LPL (Table 1; Figures 1-3). HDL-cholesterol, LDL-cholesterol and triglycerides were not statistically different within the various sub-sets of SLE (Table 2).

There was no significant difference in plaque when anti-oxLDL positive / ALPL positive or anti-oxLDL negative / ALPL negative SLE subjects were compared to either anti-oxLDL positive/ALPL positive or anti-oxLDL negative/ALPL negative normal controls.

The observation that (a) anti-oxidized LDL and plaque is significantly increased in SLE subjects with anti-Ro60 autoantibodies and (b) plaque is elevated significantly in ENA negative SLE subjects compared to normal control appears to be the most interesting result obtained from this study.

## Discussion

Studies reveal that SLE subjects have a risk for myocardial infarction ranging from 9 to 50 fold compared to that of the general population.^24^ While traditional risk factors contribute to CVD in SLE, non-traditional risk factors play an important part.^25,26^ Contrary to that seen in general population, young premenopausal lupus subjects more commonly have premature CVD.^1^

In addition, the disorder is uncharacteristic in SLE because it is not linked with the traditional inflammatory problem typical of normal atherosclerosis, like increased C-reactive protein and plasma low-density lipoprotein. SLE subjects suffering a cardiovascular event tended to more commonly have lupus diagnosis at an older age, lengthier duration of lupus, longer period of corticosteroid use, hypercholesterolemia and postmenopausal status than SLE subjects without an event.^24,25^

That immune dysregulation typical of lupus is important in plaque progression and vascular complications is borne out by the observation that higher damage index score and less aggressive immunosuppression are connected with increased CVD problem.^27,28^ Both innate and adaptive immune systems that bring about the inflammatory state of lupus may also be associated with development and progression of CVD.^29^ Results of two earlier studies from our group and data from this study support an important role for autoantibodies as a potentially increased risk for atherosclerosis in SLE.

Our study looked at the association of autoantibodies targeting RNP, SmRNP, Ro, La, oxidized LDL, LDL or lipoprotein lipase with carotid intima-media thickening in SLE and control subjects. We found that plaque, denoted by carotid intima-media thickening, was the highest in the group with anti-Ro. This group also had statistically significant high levels of antibodies targeting oxidized LDL.

Studies show that the measurement of plaque by ultrasound of the carotid arteries is a useful predictor of coronary artery disease associated with coronary disease events like angina and myocardial infarction.^30,31^

We know that patients with anti-Ro and anti-La have highly statistically significant elevations of anti-oxidized LDL and anti-phospholipid antibodies.^23^ We studied anti-oxLDL antibodies in an SLE patient over a period of 137 months. We found anti-oxLDL was very high at the time we started studying this SLE patient and it stayed highly elevated for almost the entire period.^23^ We have also found that rabbits immunized with Ro60 or Ro peptides develop an SLE-like disease wih high levels of anti-Ro60 and intermolecular epitope spreading to La, oxidized LDL and phospholipids (manuscript in preparation). However, the exact mechanism by which plaque and antibodies against oxidized LDL arise in SLE subjects with anti-Ro is not known.

Ultraviolet irradiation or caloric stress can induce increased expression of autoantigens. The ultraviolet irradiated skin is targeted better by anti-Ro antibodies under experimental conditions. This manifestation has been appreciated in the skin of subjects with sub-acute cutaneous lupus erythematosus. In these subjects the autoantigen availability increases after sun exposure and such a factor can induce *in situ* formation of Ro/anti-Ro immune complex.^32^ Ultraviolet exposure has been shown to increase free radical release by skin cells.^33^ Increased production of reactive oxygen species by this process or by the depletion of anti-oxidants or anti-oxidant enzymes by autoantibodies^34,35^ can increase oxidative stress, which can lead to LDL oxidation. Anti-lipoprotein lipase autoantibodies found in SLE is thought to allow LDL to be long lived in the circulation, by hindering lipid transport further downstream.^11^ Therefore LDL becomes an ideal candidate for oxidative modification. LDL modified in this fashion behaves like a neoantigen, allowing the host to see them as non-self antigen and inducing the host to make antibodies against such antigens.^36,37^ Free radicals and antibodies to oxidized LDL have been implicated in atherosclerosis found in SLE.^38^

Oxidized LDL has been shown to complex with plasma β2-glycoprotein I (β2GPI) and becomes autoantigenic, eliciting the production of specific antiphosholipid antibodies.^39^ We recently reported significantly elevated IgG anti-phospholipid that correlated with anti-oxidized LDL in SLE.^23^ In an interesting twist to the investigative process, the question now being asked is whether atherosclerosis is an autoimmune disease. Elevated levels of oxLDL/β2GPI were first observed in SLE and antiphospholipid syndrome (APS), and subsequently with coronary heart disease and type 2 diabetes mellitus. When subjects with chronic CHD were studied prospectively over a 2-year period, the early plasma concentrations of oxLDL/β2GPI was found to correlate with the number and severity of cardiovascular events.^39^ Recent work shows an association between anti-Ro and low IgM anti-phosphoryl choline antibodies is SLE patients. Low IgM anti-phosphoryl choline is associated with CVD in subjects that do not have anti-phospholipid antibodies.^40^

Some limitations to this study include the lack of data on organ damage. Also, we do not have data on tobacco smoking for these subjects. Hence the data was not adjusted for traditional risk factors for CVD such as smoking.

It would be of great interest to know if SCLE subjects have increased plaque and elevated levels of antibodies against oxidized LDL and lipoprotein lipase, since they do not have much systemic inflammation. It would also be of interest to investigate cardiovascular events in relation to autoantibody subsets in SLE. It is possible that cardiovascular events are augmented in SLE subset with anti-Ro and elevated anti-oxidized LDL antibodies, which will have significant clinical consequences.

## Acknowledgement

The authors are grateful for the contributions of Dr. Morris Reichlin and Marianne Reichlin, Oklahoma Medical Research Foundation. This work was supported by Grant # ARO53483 from Rheumatic Disease Research Cores Center, Oklahoma, Grant # GM104938 from Clinical and Translational Science Award, NIJ and Grant # A1082714 from ACE to RHS.

